## Supplementary for "Micro-PINGUIN: Microtiter plate-based ice nucleation detection in gallium with an infrared camera"

### Supplementary material

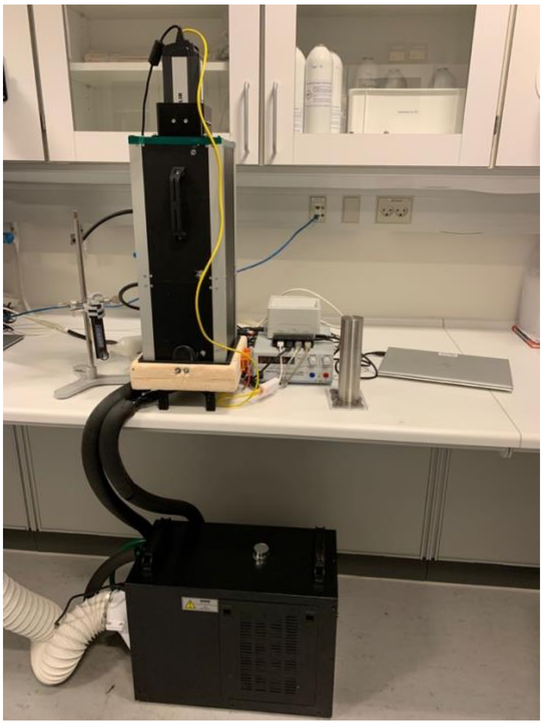

S 1: Photograph of the micro-PINGUIN instrument. The water cooling on the floor is attached to the cooling unit of the instrument. A camera tower with the infra-red camera is positioned on top of the cooling unit and records the freezing events. The weight used to mount the PCR plate in the gallium both is standing next to the computer.

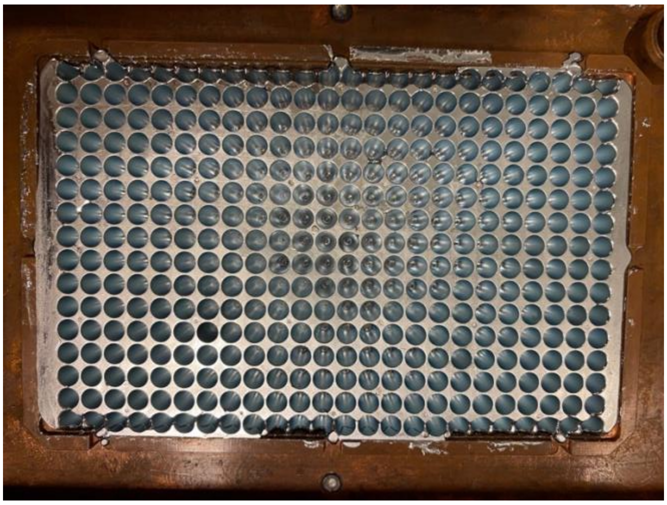

S 2: Photograph of the solid gallium after removing the PCR plate. The gallium adapts to the shape of the PCR plate during solidification.

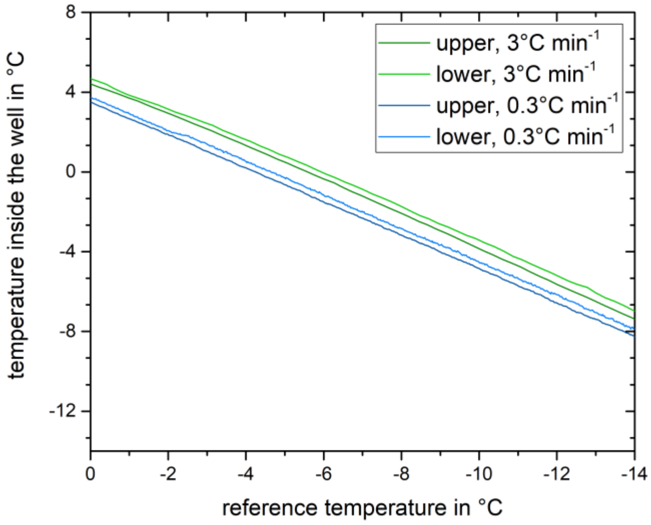


S 3: Vertical gradient measured at the top and the bottom of the central well with an external thin thermistor. The gradients are similar for the cooling rate of 0.3 °C $min^{-1}$ and 3 °C $min^{-1}$.

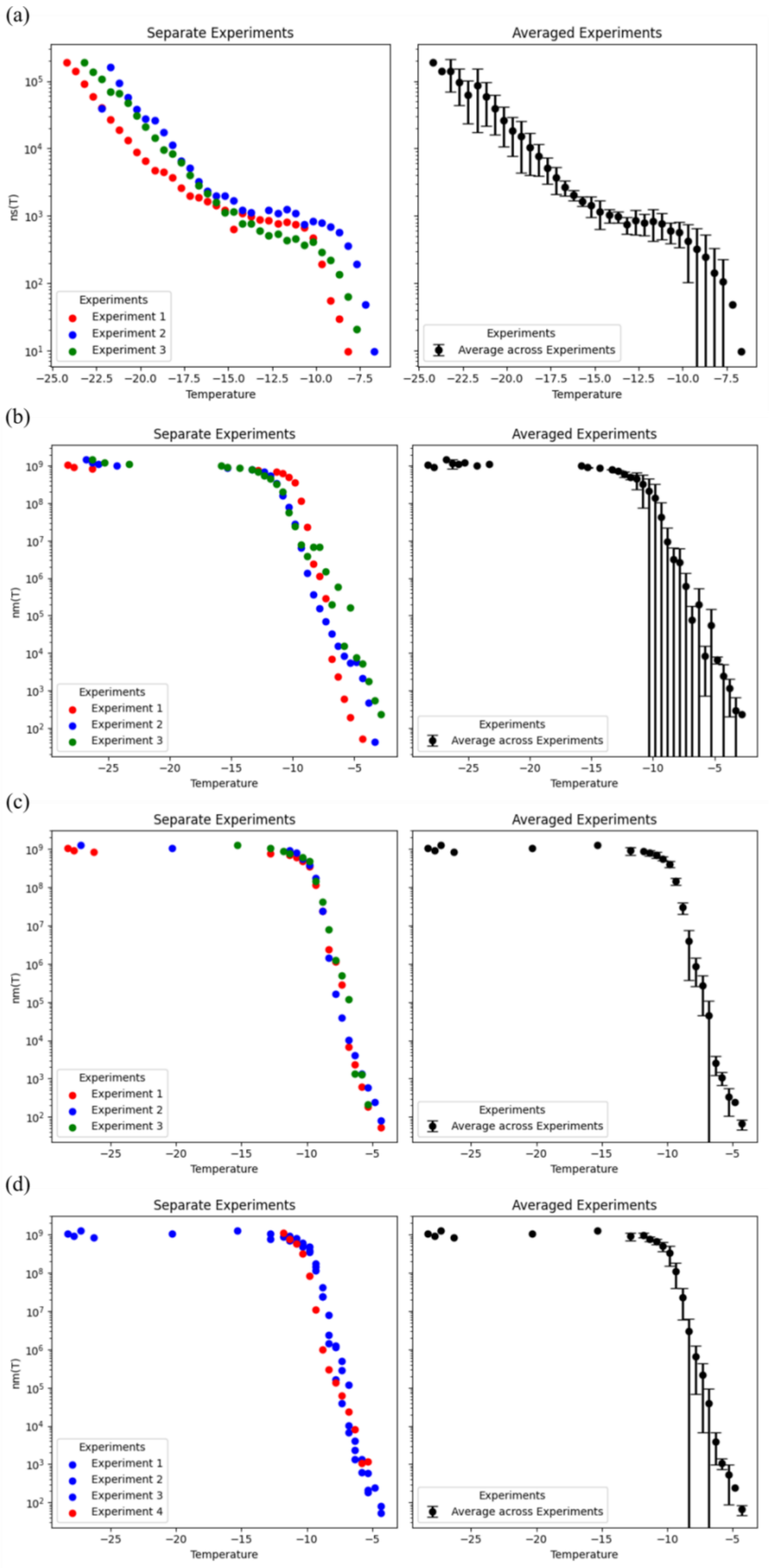


S 4: (a) Number of INPs per surface area for three Illite NX suspension measurements. Left: Data for individual experiments binned in 0.5°C temperature bins. Right: Average value and standard deviation between the three measurements. (b) – (d) same as in (a) but for the number of INPs per mg Snomax for three separately prepared suspensions (b), for three frozen aliquots from one suspension and (c) for three frozen aliquots (blue) and after 4 months storage time (red).

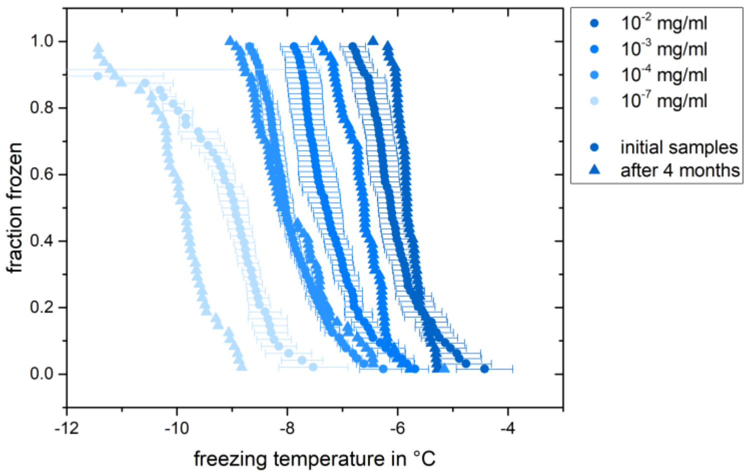


S 5: Effect of storage for the Snomax suspension. The round symbols show the average and standard deviation from three aliquots measured after 1 day storage in the freezer and the triangles show the results for one measurement after 4 months storage. The data for $10^{-5}$ mg $ml^{-1}$ and $10^{-6}$ mg $ml^{-1}$ are not shown for illustrative purposes.
